## Supplementary figures and images for "Consecutive treatments of methamphetamine promote the development of cardiac pathological symptoms in zebrafish"

### Supplemental Figure 1

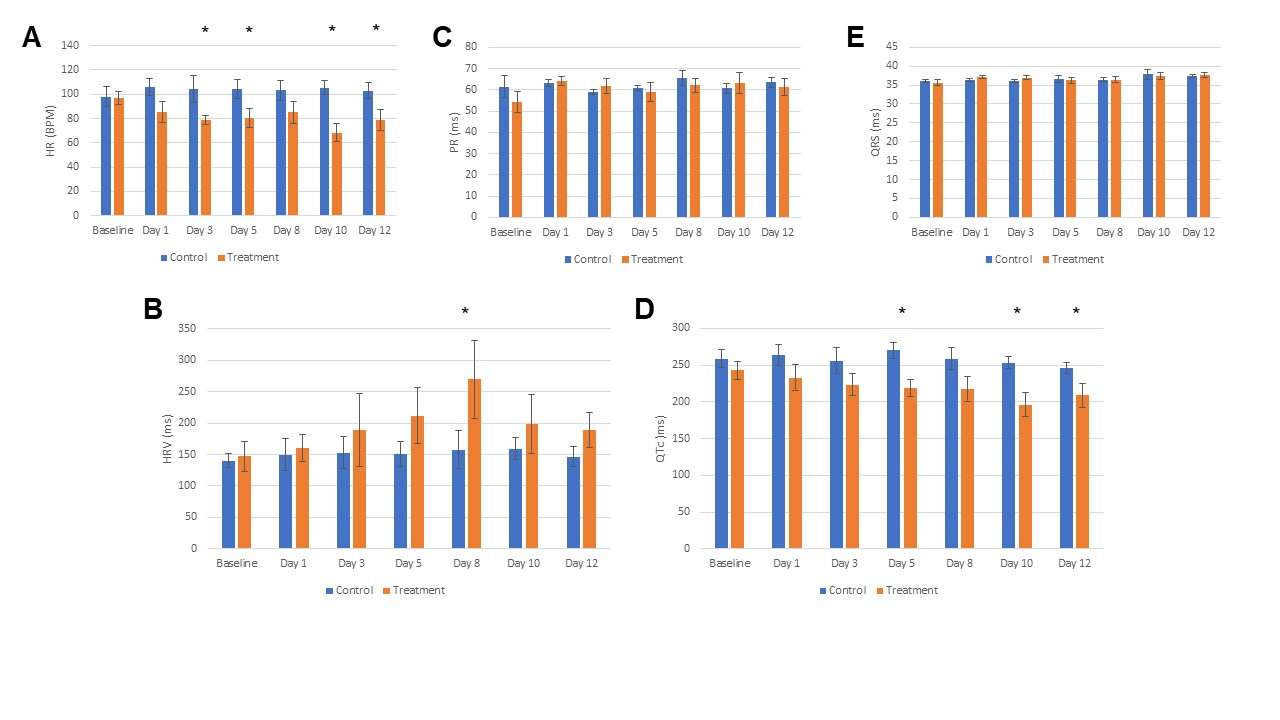
